# Ghosts of a structured past: Impacts of ancestral patterns of isolation-by-distance on divergence-time estimation

**DOI:** 10.1101/2020.03.24.005736

**Authors:** Zachary B. Hancock, Heath Blackmon

**Affiliations:** Department of Biology at Texas A&M University; Ecology & Evolutionary Biology Interdisciplinary Program at Texas A&M University

**Author notes:** Author contributions: HB conceptualized the study; ZBH performed the analyses; ZBH and HB wrote the manuscript.

**Keywords:** Phylogenetics, divergence-time, isolation-by-distance

## Abstract

Isolation by distance is a widespread pattern in nature that describes the reduction of genetic correlation between subpopulations with increased geographic distance. In the population ancestral to modern sister species, this pattern may hypothetically inflate population divergence time estimation due to the potential for allele frequency differences in subpopulations at the ends of the ancestral population. In this study, we analyze the relationship between the time to the most recent common ancestor and the population divergence time when the ancestral population model is a linear stepping-stone. Using coalescent simulations, we compare the coalescent time to the population divergence time for various ratios of the divergence time over the product of the population size and the deme number. Next, we simulate whole genomes to obtain SNPs, and use the Bayesian coalescent program SNAPP to estimate divergence times. We find that as the rate of migration between neighboring demes decreases, the coalescent time becomes significantly greater than the population divergence time when sampled from end demes. Divergence-time overestimation in SNAPP becomes severe when the divergence-to-population size ratio < 10 and migration is low. We conclude that studies estimating divergence times be cognizant of the potential ancestral population structure in an explicitly spatial context or risk dramatically overestimating the timing of population splits.

## Introduction

A major goal in phylogenetic and phylogeographic studies is the estimation of species divergence times. The topic has a long and contentious history largely centered around questions of how to appropriately apply fossil calibrations (e.g., Heath et al. 2014; Brown and Smith 2018), rate heterogeneity (Pond and Muse 2005), rate of morphological evolution (Lynch 1990), and selecting an adequate clock model (Douzery et al. 2004; Lepage et al. 2007).

Beyond methodological concerns are those that emerge from the nature of the data itself. Most phylogenetic models assume that fixed differences between species are the result of genetic drift, and under the neutral theory of molecular evolution (Kimura 1968; King and Jukes 1969) the rate of evolution (or substitution rate) is equal to the per generation neutral mutation rate, *μ* (Kimura 1983). For well-calibrated molecular clocks (e.g., Knowlton and Weigt 1998; Weir and Schluter 2008; Herman et al. 2018), we can estimate the time of divergence (usually in years) as π_12_ / 2*μ*, where π_12_ is the pairwise sequence divergence between species 1 and 2. However, in general we are not interested in estimating the divergence time of specific genetic variants, but rather the time of population divergence (T_D_). For example, we might be interested in estimating the timing of a vicariant event that we suspect corresponds to a past geological upheaval.

There is a known discrepancy between the coalescent time of neutral genetic variants (T_MRCA_) and T_D_ (Nei and Li 1979). The degree of this discrepancy is determined by the ratio of T_D_ / *N*_e_, where *N*_e_ is the effective population size (Edwards and Beerli 2000; Rosenberg and Feldman 2002). First, lineages must be within the same population, which occurs T_D_ generations in the past; second, these lineages must then coalesce, which on average requires 2*N*_e_ generations. Therefore, for a completely panmictic population: T_MRCA_ = T_D_ + 2*N*_e_. The expected amount of pairwise sequence divergence is

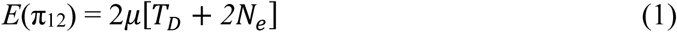

(Wakeley 2000). When the ratio of T_D_ / *N*_e_ is large, the bias in coalescent time in the ancestral population is minimal compared to T_D_ (Edwards and Beerli 2000). However, as T_D_ / *N*_e_ becomes small, 2*N*_e_ plays a major role in the overall sequence divergence between species. Nordberg and Feldman (2002) evaluated the relationship between T_MRCA_ and T_D_ in a simple two population split model using coalescent simulations. They found that T_MRCA_ converged on T_D_ when the ratio of T_D_ / *N*_e_ ≈ 5. Importantly, the *N*_e_ in these models is that of the ancestral population; therefore, the extent of overestimation is the result of demographic conditions present in the ancestor. Demographic conditions that inflate *N*_e_, such as ancestral population structure or a bottleneck following the split, is expected to have a major impact on divergence-time estimation (Gaggiotti and Excoffier 2000; Edwards and Beerli 2000; Wakeley 2000).

Wakeley (2000) demonstrated that in descendant species who share an ancestor whose population dynamics are characterized by an island model (Wright 1931) with free migration between demes, overestimation of divergence-times are on the order of 2*N*_*e*_*D*[1 + (1/2M)] where M = 2*N*_*e*_*mD*/(*D* - 1) and *m* is the migration rate. The expected amount of pairwise sequence divergence is therefore

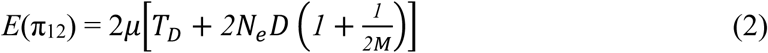

where *D* is the number of demes.

Population subdivision initially leads to shallow coalescent times where individuals within a shared deme rapidly find ancestors (the “scattering phase”). However, since ancestral lineages must be in the same deme to coalesce, the rate in the “collecting phase” is characterized by the migration rate that shuffles ancestors around the range, reducing the probability that lineages coalesce (Wakeley 1998; 1999).

In the context of real populations, the island model of migration rarely applies (Meirmans 2012). Instead, population structure is the product of the spatial distribution and dispersal potential of the organism in question. Often this structure is in the form of isolation-by-distance (IBD). IBD is a widespread pattern in natural systems, characterized by a reduction in the probability of identity by descent (Wright 1943) or genetic correlation (Malécot 1968) with geographic distance. Patterns of IBD are most pronounced in stepping-stone models (Kimura 1953; Kimura and Weiss 1964) in which migration is restricted to neighboring demes. In this way, demes in close proximity share a greater proportion of migrants than they do with more distant demes. Distributions of coalescent times in stepping-stone models have been studied both in the context of one dimensional and two-dimensional models that are circular or toroidal (Maruyama 1970a; 1970b; Slatkin 1991), and in continuous models with joined ends (Maruyama 1971) or with discrete edges (Wilkins and Wakeley 2002). Slatkin (1991), using a circular stepping-stone model, showed that the probability for two genes sampled *i* steps apart have an average coalescent time:

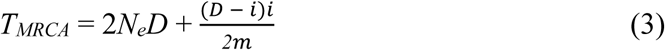

Therefore, the amount of expected pairwise sequence divergence is:

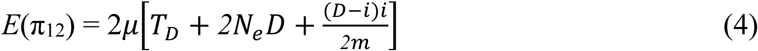

The circular stepping-stone model should overestimate T_D_ more dramatically as the number of demes becomes large and the distance between them increases. However, like the island model of free migration, circular ranges are likely rare in nature. Instead, natural populations are characterized by discrete range edges where end demes may only receive migrants from one direction (e.g., Peterson and Denno 1998; Broquet et al. 2006; Aguillon et al. 2017). Hey (1991) showed analytically in the case of a linear stepping-stone model that the distribution of coalescent times of two alleles from demes at the extremes of the range should coalesce much deeper than any two alleles chosen randomly from the population.

Ultimately, the degree to which T_MRCA_ impacts phylogenetic inference and divergence-time estimation is dependent on its impact on π_12_. Given that lower migration rates lead to greater T_MRCA_ (Hey 1991), we expect that differentiation (π_12_) between end demes compared to central will become more pronounced at smaller *m*. If the difference between the T_MRCA_ of central demes and end demes is dramatic enough, we expect that divergence dating of species that arose from ancestral end demes may significantly overestimate T_D_. ##examples

In this study, we estimate mean T_MRCA_ for two genes sampled in descendant species (either from the ends or the center of the ancestral range) in which the ancestral population is characterized by a stepping-stone model with discrete ends using a simulation approach. In particular, we are interested in what value of T_D_ / *ND* we expect T_MRCA_ to converge on T_D_. Next, we examine the distribution of π_12_ across the genome under different simulated migration conditions to compare with expectations under a panmictic model. Finally, we test the performance of the phylogenetic inference program SNAPP (Bryant et al. 2012) on simulated single nucleotide polymorphism (SNP) data to evaluate how these trends may bias our inference of species divergence times.

## Methods

### Coalescent simulations

Using *fastsimcoal2* (Excoffier et al. 2013), we simulated sister species with a shared ancestor whose population dynamics are characterized by a stepping-stone model. Specifically, each simulation consisted of 10 demes (*D*) with no shared migration between them until time T_D_. At T_D_, migration resumes between demes in a linear stepping-stone fashion. In *fastsimcoal2*, the migration rate is the probability of an individual from deme *i* migrating to deme *j*, where *i* and *j* are neighboring demes. Center demes receive migrants from neighboring demes at rate 2*m*, whereas demes at the end of the range receive migrants at rate *m*. This is due to the fact that end demes have only a single neighbor, whereas all center demes have two neighbors (Fig. 1A). Throughout, we will use “end demes” to represent species descending from the ends of the ancestral range; “center demes” are those that descend from the center. We sampled *k* = 2 individuals to coalesce—in one run, we sample the end demes, and in the following we sample central neighboring demes. This was performed for migration rates of 0.1, 0.01, and 0.001, and a range of T_D_ / *ND* values. In addition, we simulated an island model of migration for comparison with the stepping-stone model. In the island model, the ancestral population consisted of 10 demes with free migration between each at rate *m*. This resulted in a total of 84 distinct simulation scenarios, and each were replicated 1,000 times.

**Figure 1.**
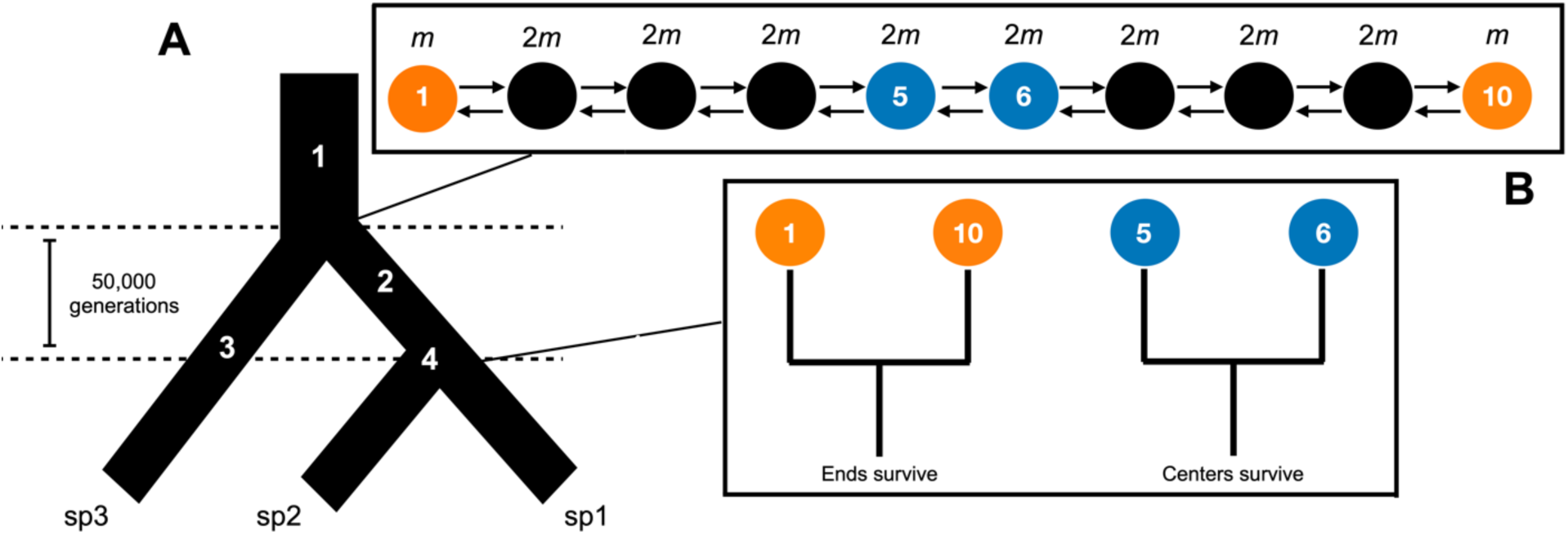
Population model for simulations. **A)** Three-taxon species tree: 1) coalescent simulations in *msprime* with *N* = 2000; 2) ancestral stepping-stone conditions begin (see **B**); 3) *N* = 1000, panmictic; 4) population split, leaving end or center demes surviving as sp1 and sp2. **B)** Ancestral population dynamics. Orange circles are “end demes” and blue circles are sampled “center” demes.

To statistically compare between the three models (end deme sampled in stepping-stone, center deme in stepping-stone, and the island model), we subset ratios of T_D_ / *ND* to values of 10, 5, 2, 1, 0.5, and 0.1. Resulting T_MRCA_ distributions for each population model were compared using a pairwise Wilcoxon test in the R platform (R Core Team 2019), as the resulting distributions were non-normal.

### Genome simulations

To evaluate how ancestral IBD impacts pairwise sequence divergence (π_12_), genome-wide coalescent times (T_MRCA_), and divergence-time estimation, we performed hybrid simulations that combined the coalescent simulator *msprime* (Kelleher et al. 2016) and the forward-time simulator SLiM v3.3 (Haller and Messer 2019). Since forward-time simulators begin with individuals that are completely unrelated, often a neutral burn-in period is required to allow coalescence or mutation-drift equilibrium to occur (Haller et al. 2019). This can be computationally costly and time consuming; however, using tree-sequence recording methods in SLiM (Haller et al. 2019) we can bypass the need to equilibrate during the forward-time simulation. To generate a panmictic ancestral population with a coalescent history, we simulated 2000 individuals (*N*_e_ = 4000) using *msprime* with genome sizes of 10 Mb and a recombination rate of 10^−8^ (∼0.1 recombination events per individual per generation). The resulting coalescent trees were then imported into SLiM as the basis for the starting population.

In SLiM, the initial population was split into two populations of *N* = 1000: 1) an outgroup that remained panmictic (“sp3” in Fig. 1A) and 2) the ancestral population, which was subdivided into 10 demes (*N* = 100 per deme) in a linear stepping-stone model. These dynamics persisted for 50,000 generations after which the ancestral population was split into either “end” demes or “center” demes (see Fig. 1A). Population sizes of each deme following the split was increased to 1000 to maintain *N* throughout the simulation. Five different T_D_ values were simulated which correspond to T_D_ / *ND* ratios of 50, 25, 10, 5, and 1. These values of T_D_ / *ND* were chosen based on the results from the coalescent simulations (see Results); for values >10, T_MRCA_ is expected to converge on T_D_, whereas values <10 are expected to overestimate T_D_ regardless of migration rates.

The resulting tree-sequences from the SLiM simulation were imported into Python 3 using *pyslim*, and we overlaid neutral mutations (*μ* = 10^−7^) onto the trees using *msprime*. Pairwise divergence (π_12_) was then estimated across the genome in windows of 100 kb for both end demes and center demes. These values were also converted into generations using π_12_ / 2*μ*, which gives a rough estimate of divergence time per window.

By rearranging equation 1, we can naively calculate *N*_e_ for the ancestral population from genome-wide π_12_ as:

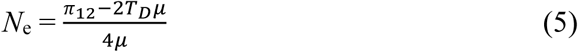

From this, we plot estimated ancestral *N*_e_ within 100 kb windows across the genome to compare with the known census population size (*N*_c_ = 1000), and to evaluate the relationship between *N*_e_ and *N*_c_ in the presence of IBD.

Next, we plotted the distribution of coalescent times (T_MRCA_) across the genome to visualize differences between T_MRCA_ of end and center demes. Median T_MRCA_ for each ratio and migration rate was compared via a Kruskal-Wallis test and a pairwise Wilcoxon rank test in R due to the data violating normality.

Each simulation produced >200,000 SNPs. For divergence-time analysis, we randomly sampled 3000 SNPs—a number found by Strange et al. (2018) to optimally perform in SNAPP (Bryant et al. 2012). Each run consisted of 10 individuals from species sp1 and sp2, and 1 individual from the outgroup population, sp3 (Fig. 1). Unlike other fully coalescent models, SNAPP does not sample from gene trees directly to estimate the species tree, but instead integrates over all possible gene trees using biallelic SNPs. The method has been found previously to perform well on both simulated and empirical data (Bryant et al. 2012; Strange et al. 2018). We designated a gamma-distributed prior on *θ* (=4*N*_*e*_*μ*) with a mean equal to the expected π_12_ (equation 1). Forward (*u*) and backward (*v*) mutation rates were estimated within BEAUti (Bouckaert et al. 2014) from the empirical SNP matrix using the tab *Calc_mutation_rates*, and these values were sampled during the MCMC. The rate parameter λ, which is the birth-rate on the Yule tree prior, was gamma-distributed with α = 2 and β = 200, where the mean is α / β (Leaché and Bouckaert 2018).

SNAPP is designed to handle incomplete lineage sorting (ILS), but to minimize its effects—since we are not interested in the program’s ability to estimate topology but rather branch-lengths—we applied a fixed species tree. Branch-lengths in SNAPP do not scale to time, but instead are measured in number of substitutions. Given a fixed mutation rate, we convert the number of substitutions separating sp1 and sp2 to the number of generations as *g* = *s* / *μ*, where *s* is branch-lengths in units of substitutions (Bouckaert and Bryant 2015). The MCMC chain length was 10–50 million sampling every 1000 with a burn-in of 10%, ensuring that ESS values of interest were all >200. Runs were performed on the high-performance computing cluster CIPRES (www.phylo.org; Miller et al. 2010).

MCMC log files were then downloaded and analyzed in R. The performance of SNAPP was evaluated by comparing traces of end and center demes across migration rates and T_D_ / *ND* values. Results were evaluated using a two-way ANOVA followed by Tukey’s HSD post hoc test in R. Trees from the MCMC were summarized in TreeAnnotator v.2.6.0 (Bouckaert et al. 2014) and visualized in R using the package *ggtree* (Yu et al. 2017). Branch colors and widths were scaled by estimated median *θ* per branch.

## Results

### Coalescent simulation results

The coalescent simulations produced trends superficially similar to those found by Rosenberg and Feldman (2002). At the lowest T_D_ / *ND*, the proportion of deep coalescence was dramatically greater than at higher values with the curve producing a similar logarithmic relationship (Fig. 2). However, T_D_ and T_MRCA_ did not necessarily converge when T_D_ / *ND* = 5. Instead, the rate of convergence was dependent on both the deme sampled and the migration rate.

**Figure 2.**
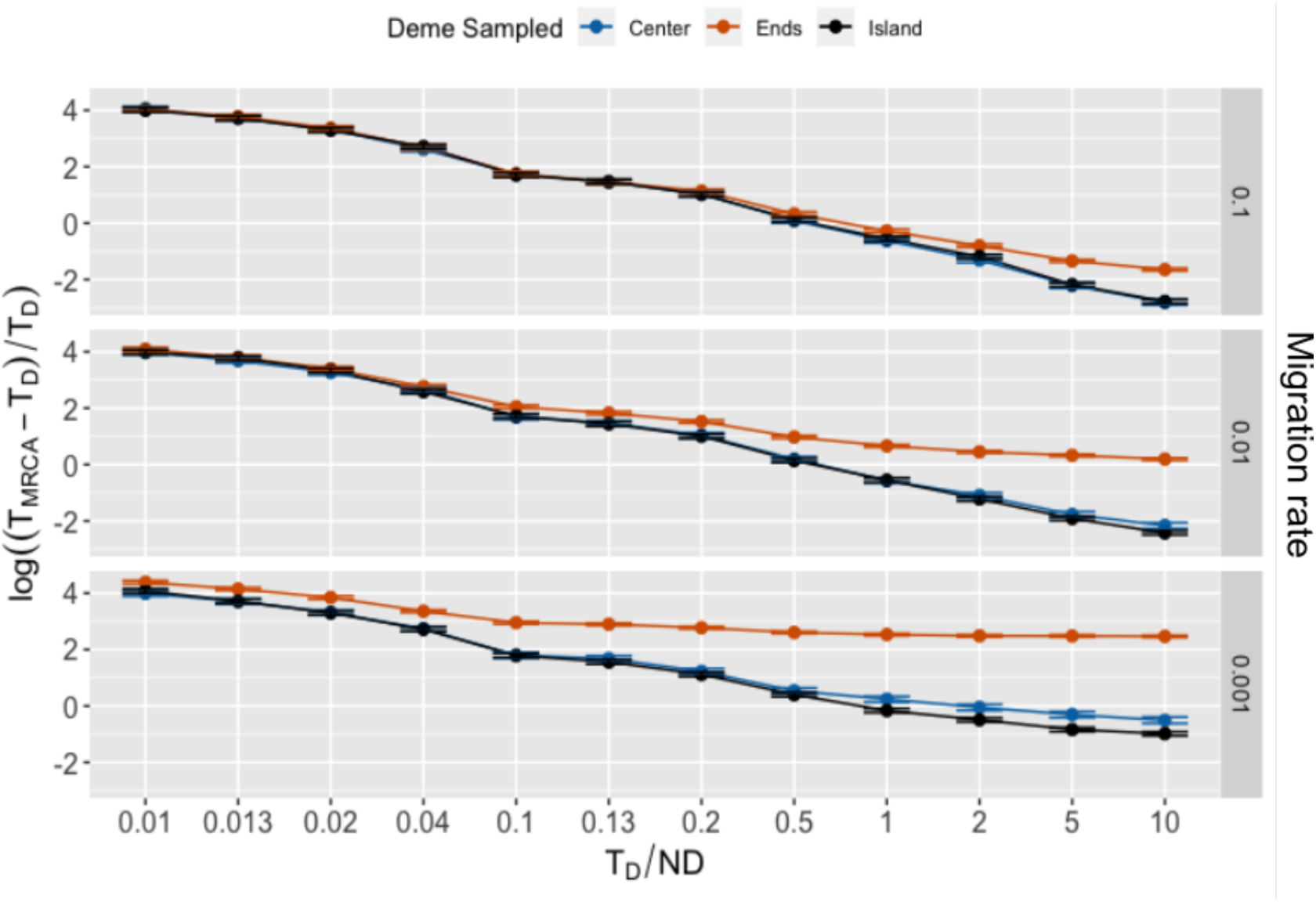
Plots of log((T_MRCA_ – T_D_) / T_D_) against T_D_ / *ND* for each migration rate (0.1, 0.01, 0.001). Each point is the mean of 1000 simulations. Y-axis has been log-transformed to aid in visualizing differences between model/deme sampled.

When migration was high (*m* = 0.1) and T_D_ / *ND* was less than 0.5, there was no significant difference between center or end demes in the stepping-stone model or the island model. However, for values of T_D_ / *ND* > 0.5, the T_MRCA_ of end demes became significantly different from both island (*p* < 0.02) and center demes (*p* < 0.01; see Table S1). When migration was reduced below 0.1, this pattern became more extreme. End demes were significantly different in all pairwise comparisons of models (*p* < 0.000001), and center demes differed from the island model at T_D_ / *ND* ratios of 0.5, 2, and 10 (*p* < 0.03) when *m* = 0.01. At the lowest migration rate simulated (*m* = 0.001), all pairwise model comparisons were significantly different when T_D_ / *ND* > 0.5 (*p* < 0.001; see Table S1).

### Genome simulation results

Results from the genome simulation approach corroborated those found with *fastsimcoal2*. Regardless of T_D_ / *ND*, when *m* = 0.1 the difference between center and end demes was less severe and only marginally significant (*p* = 0.001) relative to when *m* < 0.1 (Table S2). Across the simulated genomes, T_MRCA_ became dramatically deeper between end than center demes as migration fell below 0.01. For the genome-wide divergence estimates, the degree of overestimation depended on the ratio of T_D_ / *ND*. While all scenarios where *m* = 0.001 overestimated the true T_D_, when T_D_ / *ND* < 10 end demes were 5–60 times more diverged than expected (Fig. 3). This is a direct result of the deeper coalescent times between end demes when *m* < 0.1, as these longer branches provide more time for mutations to occur and accumulate (Fig. 4).

**Figure 3.**
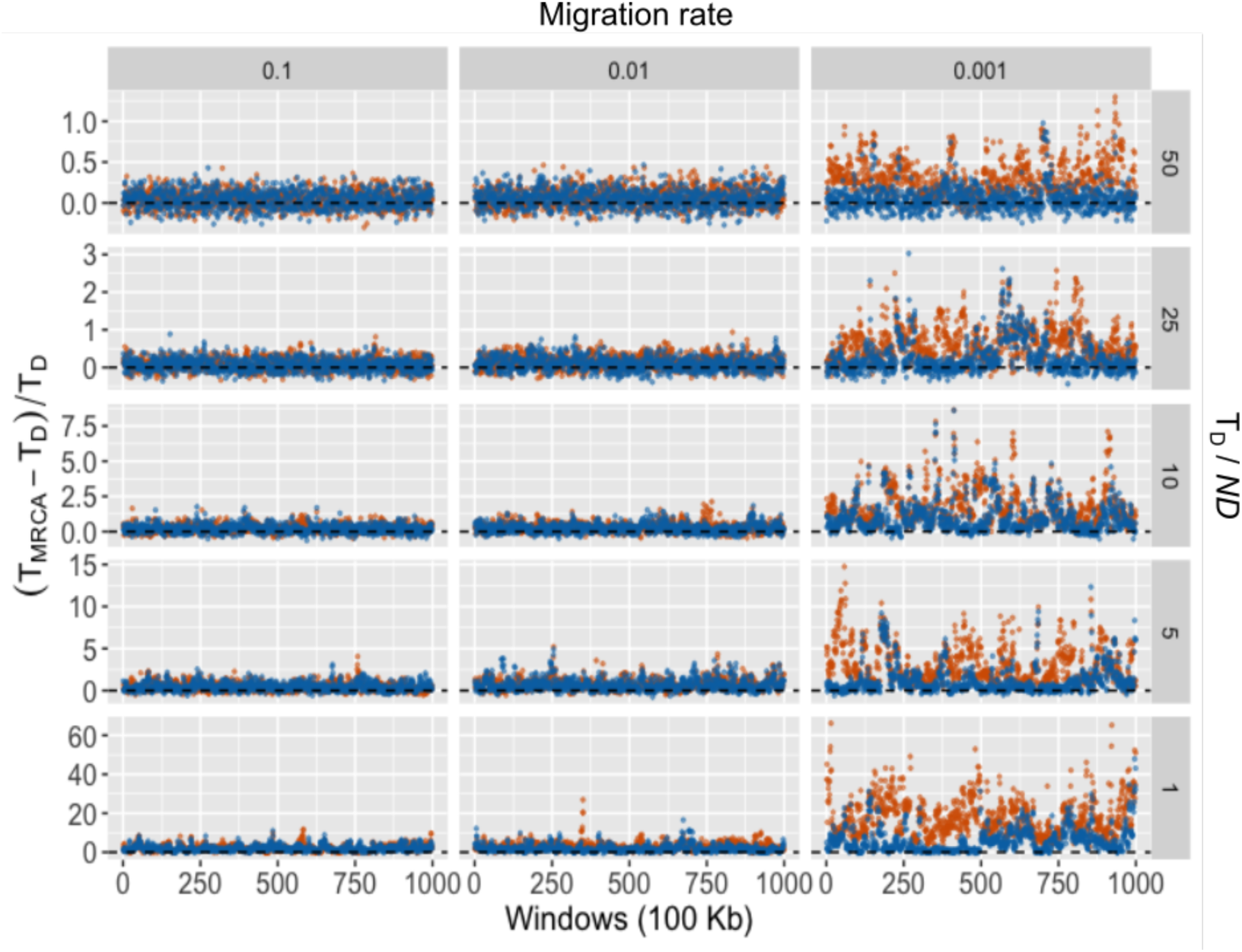
Genome-wide divergence times based on π_12_. Divergence times are estimated as π_12_ / 2*μ* and evaluated in 100 Kb windows. The y-axis is the scaled proportion of overestimation, where T_MRCA_ is the estimated age and T_D_ is the true age. The dashed line represents the value at which these two converge (i.e., 0). Center (blue), ends (orange). Note the y-axis differs between the panels.

**Figure 4.**
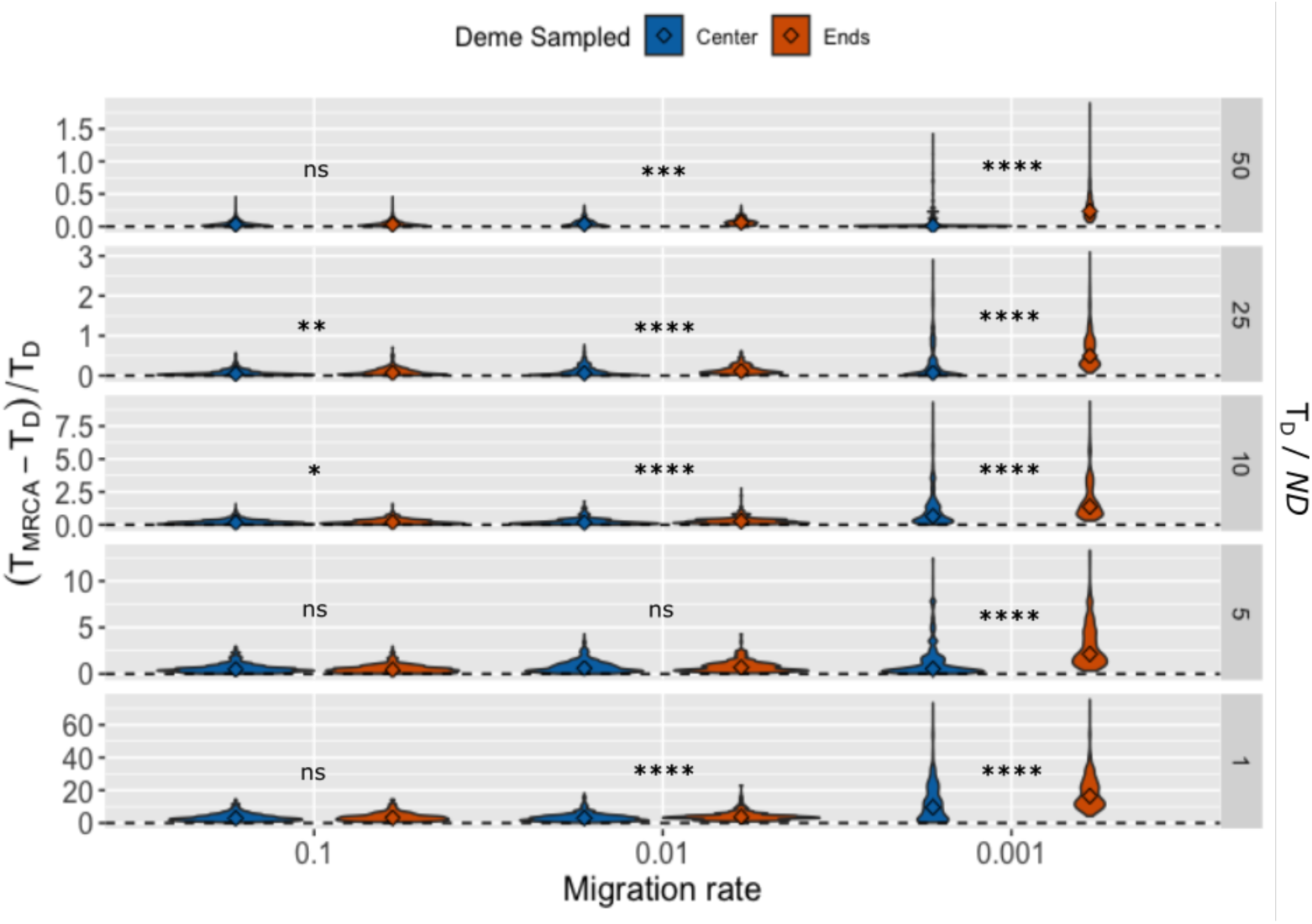
Violin plot of coalescent times (T_MRCA_) across the genome, where times have been converted into proportions of the population divergence time (T_D_). Diamonds are medians; ns = “not significant”, *p* < 0.05 (*), *p* < 0.001 (**), *p* < 0.0001 (***), *p* < 0.00001 (****). See Table S2 for specific *p*-values. Note that the y-axis differs between panels.

Genome-wide coalescent times (T_MRCA_) are shown in Fig. 4. When *m* = 0.1, only T_D_ / *ND* = 25 and 10 were significantly different between end and center demes (*p* < 0.005). Regardless of T_D_ / *ND*, the variance in T_MRCA_ steadily increased with decreasing *m*. Indeed, the increase in mean T_MRCA_ when *m* = 0.001 appears largely driven by an increase in the variance at this lower rate. Due to this, we find that ancestral *N*_e_ dramatically exceeds *N*_c_ when *m* = 0.001 (Fig. 6).

**Figure 5.**
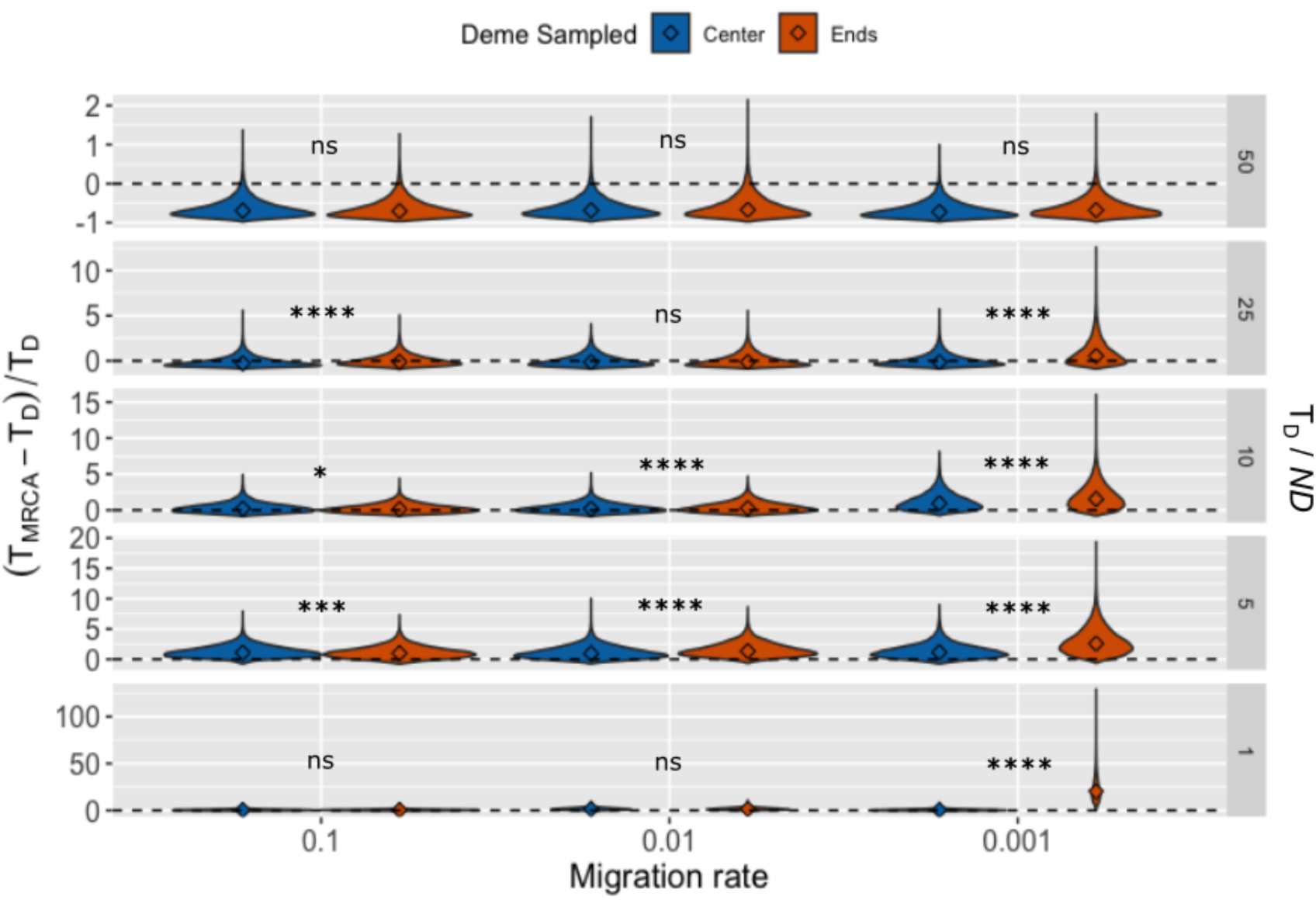
Violin plots of the estimated T_MRCA_ by SNAPP. Diamonds are medians; ns = “not significant”, *p* < 0.05 (*), *p* < 0.001 (**), *p* < 0.0001 (***), *p* < 0.00001 (****). Dashed lines represent when the estimated age converges on the true age (i.e., at 0). Note that the y-axis is different between the panels.

**Figure 6.**
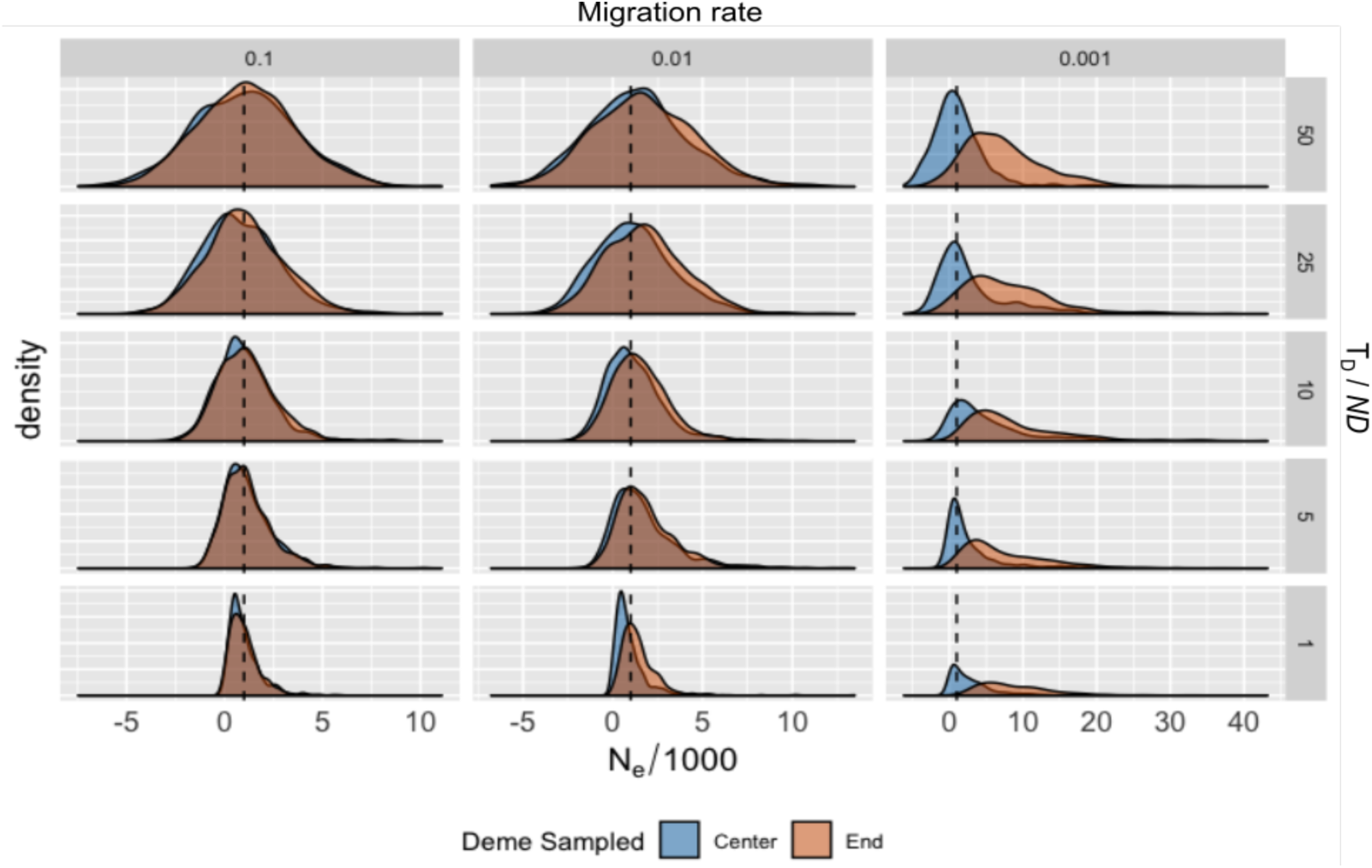
Density plot of scaled *N*_e_ (/1000) based on mean π_12_ across genomic windows of 100kb. Dashed line is when *N*_e_ / *N*_c_ = 1.

Despite the potential for divergence-time overestimation to be extreme, SNAPP was relatively resilient when T_D_ / *ND* > 10 and when *m* > 0.001. When T_D_ / *ND* = 50, SNAPP was overly conservative and underestimated the number of substitutions expected to occur (Fig. 5). When T_D_ / *ND* = 25, the mean estimate of both center and end demes when *m* > 0.001 either underestimated the true age or was within 5%. However, for end demes where *m* = 0.001 the estimated divergence time exceeded the true age by ∼80% (Table S3). A similar trend occurred when T_D_ / *ND* = 10 and 5. Here, both center and ends overestimated the true age, but the end demes did so more dramatically (138% the true age versus 81% for 10; 184% versus 67% for 5). The most dramatic overestimation occurred between end demes when T_D_ / *ND* = 1 at ∼700% the true age. Importantly, this was not merely the result of a low T_D_ / *ND* ratio, as the other migration regimes performed well. In fact, most were closer to the true T_D_ than the expected π_12_ accounting for 2*N* (Table S3).

Estimated *θ* for each branch is shown in Fig. 7 for T_D_ / *ND* = 10, and in Figs. S1–S4 for the remaining ratios. For all T_D_ / *ND* values except 1, the median ancestral *θ* was higher for end demes than center when *m* = 0.001, and the estimated *θ* for the descendant species (sp1 and sp2 in Fig. 1) was considerably lower than for the ancestor or the outgroup, sp3 (Fig. 7; Figs. S1–S3). These patterns are consistent with a population bottleneck, despite *N* being maintained throughout the simulation.

**Figure 7.**
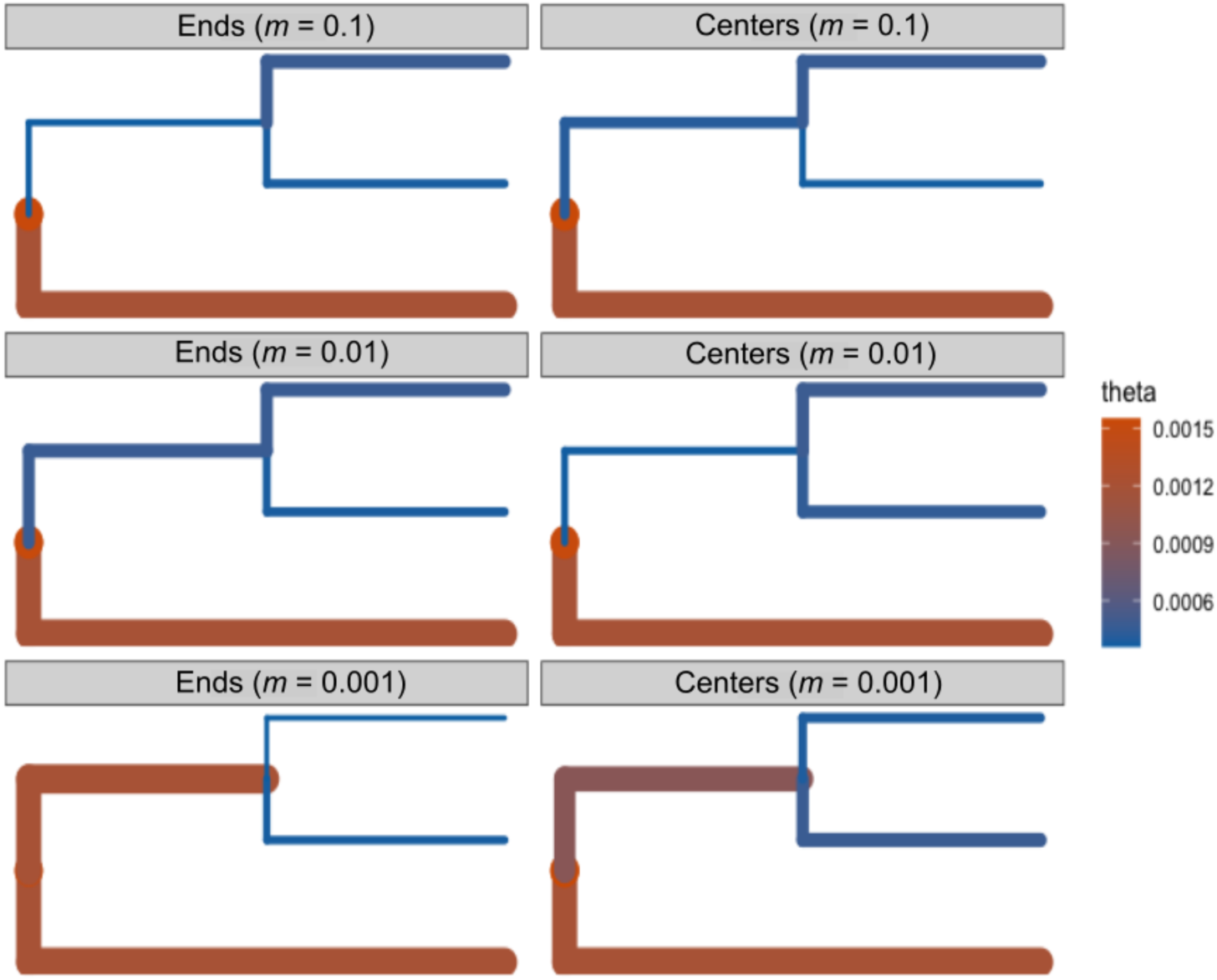
Estimates of *θ* in SNAPP for T_D_ / *ND* = 10. Branch widths are proportional to the estimated *θ*.

## Discussion

Macroevolutionary patterns are ultimately governed by microevolutionary processes (Li et al. 2018), an observation Lynch (2007), extending Dobzhansky’s (1973) maxim, summed up as “nothing in evolution makes sense except in light of population genetics”. In this light, we have demonstrated that the population genetic environment of the ancestor shapes the genetic landscape of descendant species. This has been known to impact tree topology when ILS is common (Kubatko and Degnan 2007) and overestimate divergence times in the presence of population structure caused by an island model of migration (Edwards and Beerli 2000; Wakeley 2000). Extensive prior work has shown that the stepping-stone model of migration reduces genetic correlation between demes (Kimura and Weiss 1964; Maruyama 1970a) and that demes farther apart should coalesce deeper in time than those geographically closer (Slatkin 1991; Hey 1991). However, to our knowledge, the impact of ancestral IBD has not been evaluated in the context of divergence-time estimation previously.

Rosenberg & Feldman (2002) found previously that when T_D_ / *N* = 5, T_MRCA_ and T_D_ largely converged in a simple population split model. However, when in the presence of ancestral IBD we found that convergence was dependent on the migration rate (i.e., the strength of ancestral IBD) and whether surviving demes neighbored each other or were at the range ends in the ancestral population.

When T_D_ / *ND* > 10, the ancestral dynamics contribute little to the divergence-time estimate differences between center and end demes. However, as this ratio decreases the contribution of 2*N*_e_ to overall sequence divergence becomes non-trivial. The probability that genetic variants share an ancestor just prior to the population split is higher between demes that are geographically closer than those more distant. This is mediated by the migration rate, which, when high enough, can largely erase the differences between center and end demes. When migration is high (10%, or *m* = 0.1), individuals move well between demes and the coalescent times largely converge (though deeper in time depending on the ratio of T_D_ / *ND*). However, as *m* falls below 1% (*m* = 0.01), or less than one migrant per generation being shared between demes, dispersal cannot keep up with genetic differentiation. Despite all migration regimes producing similar patterns of IBD (Fig. S5), *F*_ST_ becomes dramatically higher as migration drops below 1%. This differentiation in the ancestor contributes to the overall sequence divergence (π_12_) between species, which drives an overestimation of the time of the population split (T_D_) when end demes are the surviving lineages.

As expected, ancestral IBD skews π_12_ and T_MRCA_ away from expected values in a panmictic population, and this caused an inflation in *N*_e_ relative to *N*_c_. For T_D_ / *ND* = 50 and *m* = 0.1, the mean π_12_ for end demes was 0.010459 and 0.010419 for center demes. Using equation 5, *N*_e_ = 1147.5 for end demes and 1047.5 for central. However, when *m* = 0.001, π_12_ for end demes was 0.012948, an *N*_e_ = 7370. Center demes, on the other hand, only increased to *N*_e_ = 1255. As with the coalescent times, at lower migration rates the variance in *N*_e_ becomes exceedingly large, driving up the mean. Importantly, mean genome-wide *N*_e_ always exceeds *N*_c_ in the presence of ancestral IBD at a level dictated by the migration rate.

This feature of ancestral IBD has important consequences for conservation genetics. Many studies use *N*_e_ as a rough biological measure of population size (Turner et al. 2002; Rieman and Allendorf, 2011; Hare et al. 2011), and therefore a metric of the health of a population. However, a common phenomenon in range contractions is fragmentation and isolation (Ceballos et al. 2017), which may result in IBD. If many of the demes once contributing to the connectivity of the population have become extinct, and *N*_e_ is estimated based on the surviving demes, it will overestimate the actual number of individuals within the population (i.e., the census size, *N*_c_). Thus, we might incorrectly conclude that a population has a larger population size than it actually does, which may lead to mismanagement.

Since *N*_e_ is inflated in the ancestral lineage, the descendant species appear to pass through a bottleneck despite *N* remaining constant (Fig. 7). Estimated *θ* in SNAPP captured this dynamic with more extreme differences in *θ* (i.e., more dramatic bottlenecks) being inferred between end demes and when *m* = 0.001. Population bottlenecks have been found to cause divergence-time overestimation due to random differential survival of ancestral alleles into the descendant species (Gaggiotti and Excoffier 2000). In the presence of IBD, this differential allelic persistence between demes is mimicking a bottleneck—when demes are far apart this pattern is more extreme as they already maintain different allelic patterns ancestrally. However, because this pattern is recognizable (Fig. 7; Figs. S1–S3) it can be used to signal when ancestral IBD may be impacting our divergence-time estimation. Unfortunately, without prior range-size knowledge it may be impossible to differentiate between ancestral IBD and a bottleneck since these produce virtually identical genetic patterns. However, it may not be necessary to do so for simple divergence estimates.

The broader impact of ancestral IBD on divergence-time estimation when in the context of large phylogenies is beyond the scope of this work, but it is conceivable that the longer than expected branches between sister species might bias rate estimation (Aris-Brosou and Excoffier 1996; Magallón 2010). In the case of ancestral IBD, the inflated *N*_e_ is mimicking a pattern of substitution rate increase. Under neutrality, the rate of substitution is equal to the per generation mutation rate, *μ* (Kimura 1983); however, in the presence of population structure, substitutions may occur in the ancestral lineages between demes separated by large geographic distances. If the true age of the sister taxa is known but ancestral structure is not accounted for, the substitution rate will be upwardly biased.

Ancestral structured populations leave their imprint on descendent species in the form of greater coalescent times, and therefore larger than expected pairwise divergences between species. Further, these patterns cause inflated *N*_e_ relative to census sizes. Since ancestral IBD mimics the signature of a population bottleneck, coalescent methods that co-estimate *θ* along with the topology and π_12_, such as SNAPP and *BEAST (Bouckaert et al., 2014), may be the best suited to reveal this potential source of bias. However, fully coalescent models such as these are infamously computationally costly and not presently used for whole-genome sequence data or for phylogenies with large numbers of tips. Indeed, SNAPP becomes prohibitively slow when the number of tips is ∼30 (Leaché and Bouckaert, 2018).

In the context of larger phylogenies or organisms in which little is known about their ancestral range, it may be impossible to know if extant species descend from range centers or ends, or the level of IBD present in the ancestor. The genetic consequences of ancestral structure therefore behave much like “ghost” populations (Slatkin 2005); despite being extinct, their influence haunts our ability to adequately assess the phylogenetic history of their descendants.

## Acknowledgements

Thanks to Ben Haller and Wesley Brashear for coding help.

## Data accessibility statement

SLiM recipes, R and python code, and .XML files have been uploaded to https://github.com/hancockzb/ancestralIBD.

## Supplementary Material

**Table S1.**
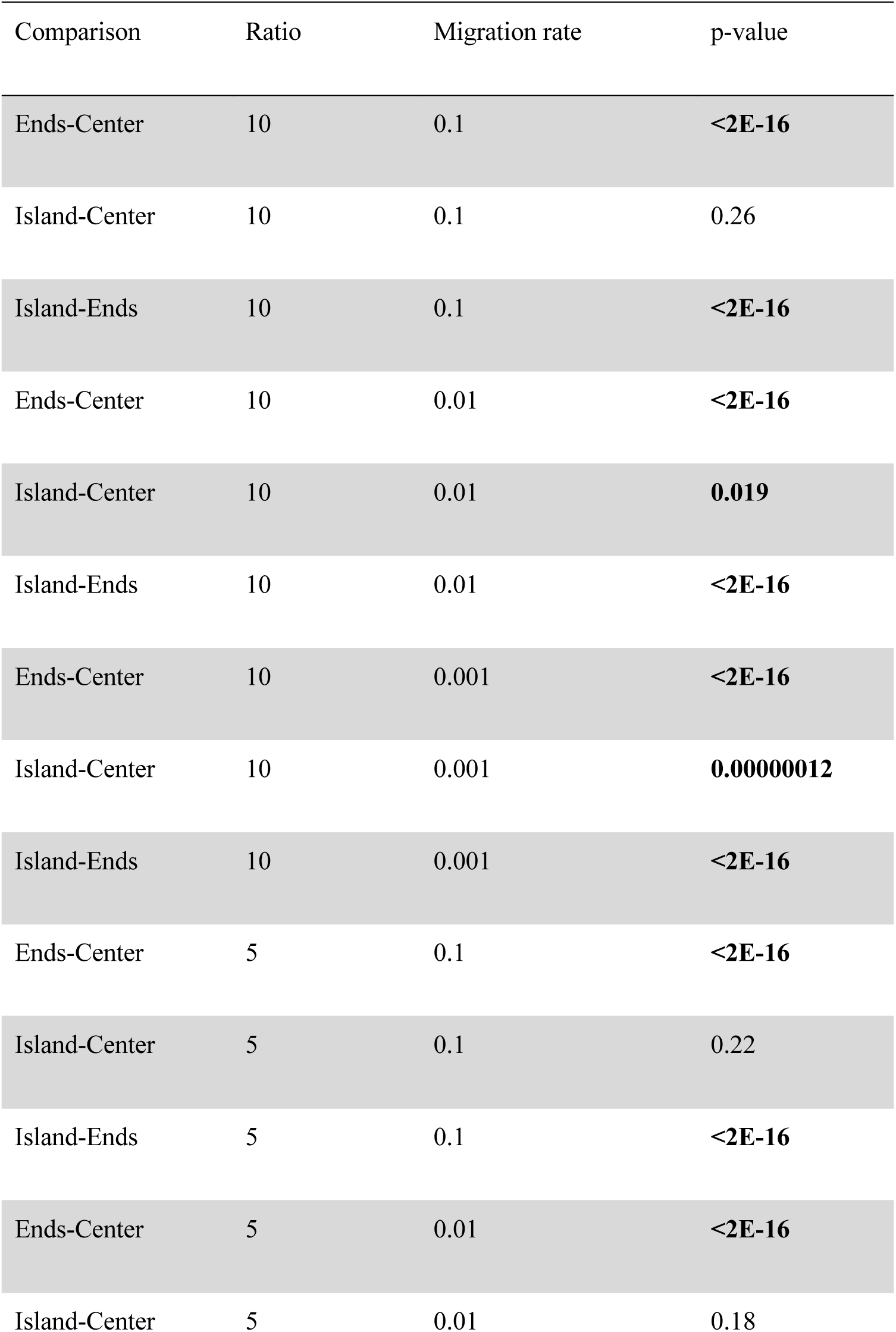
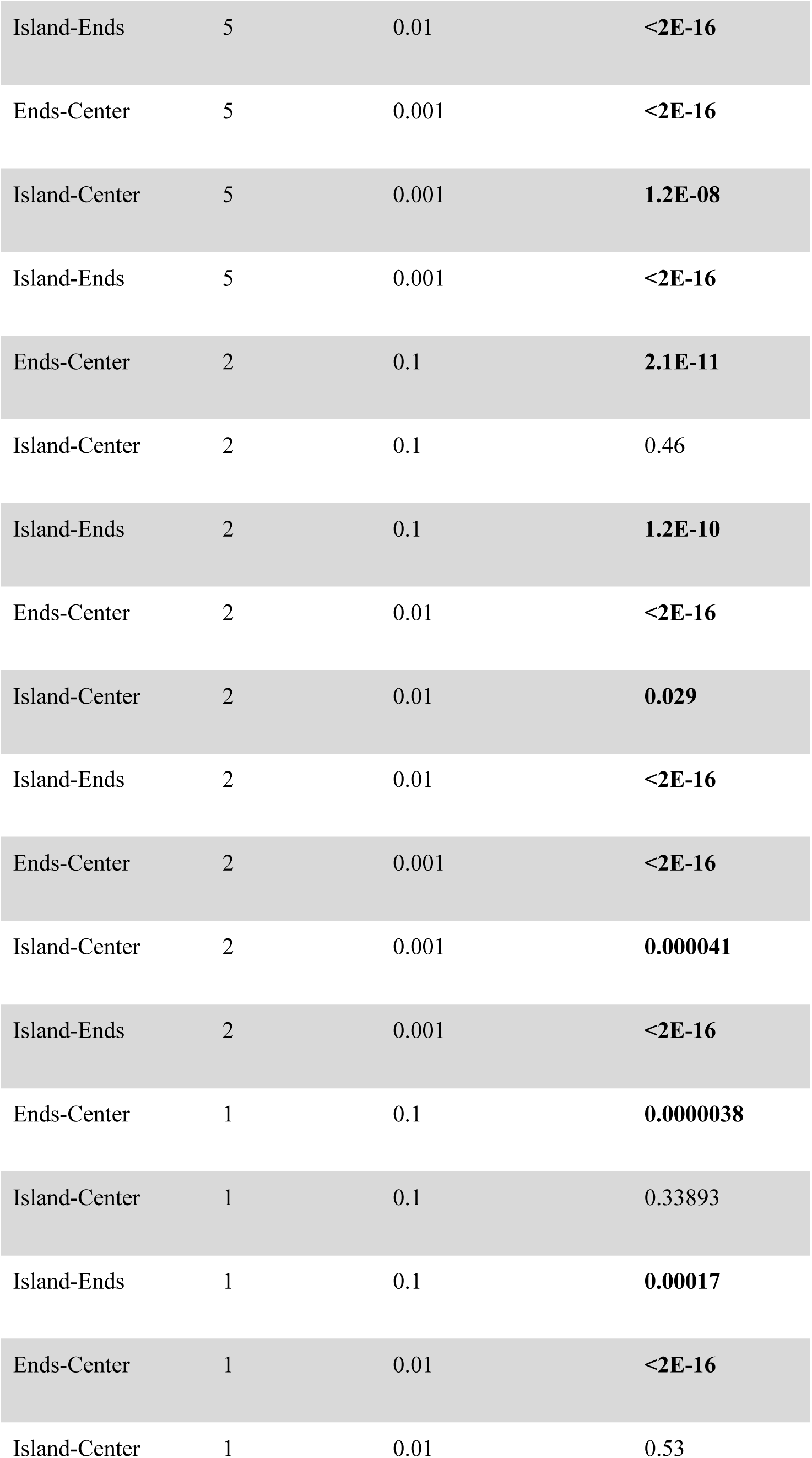
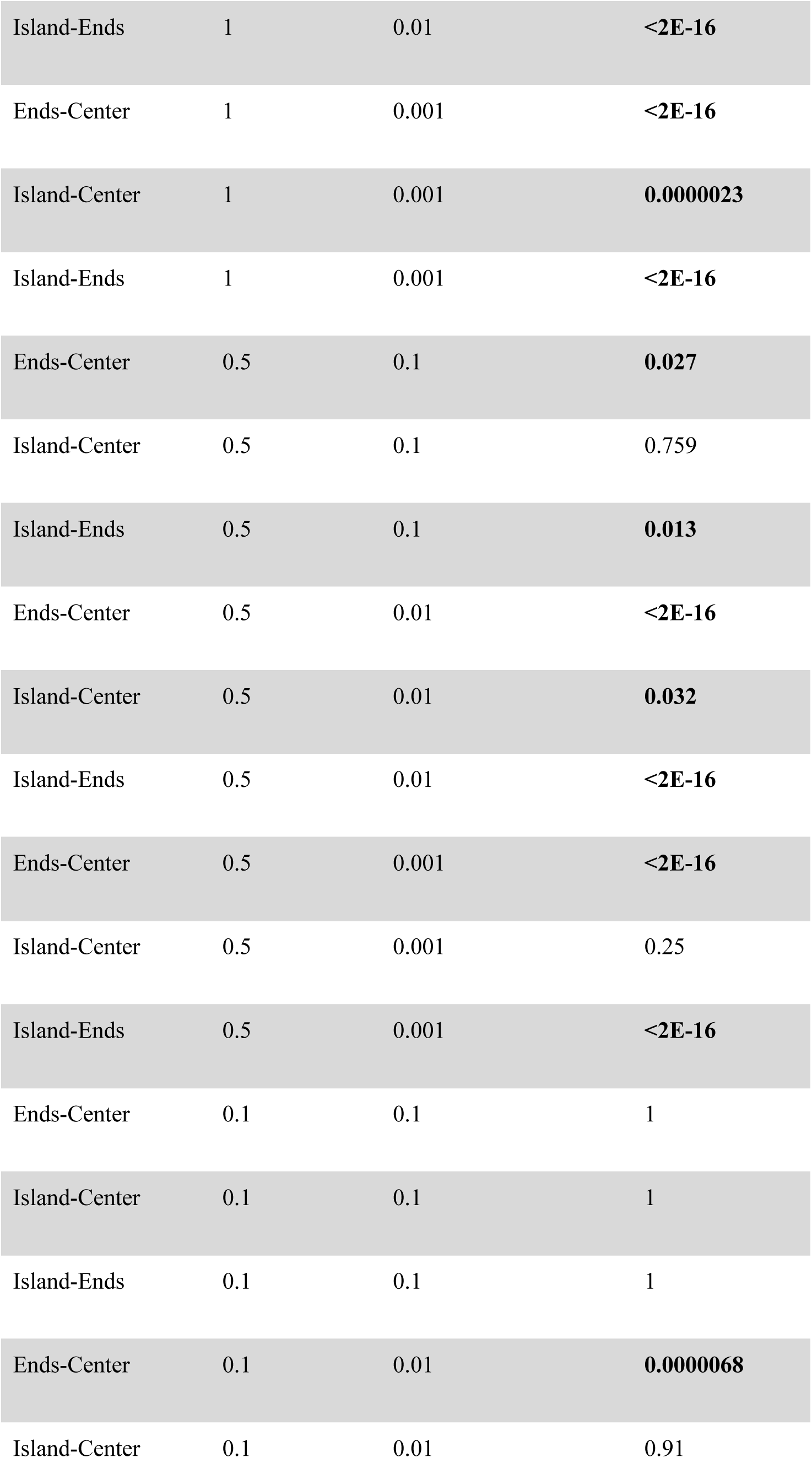
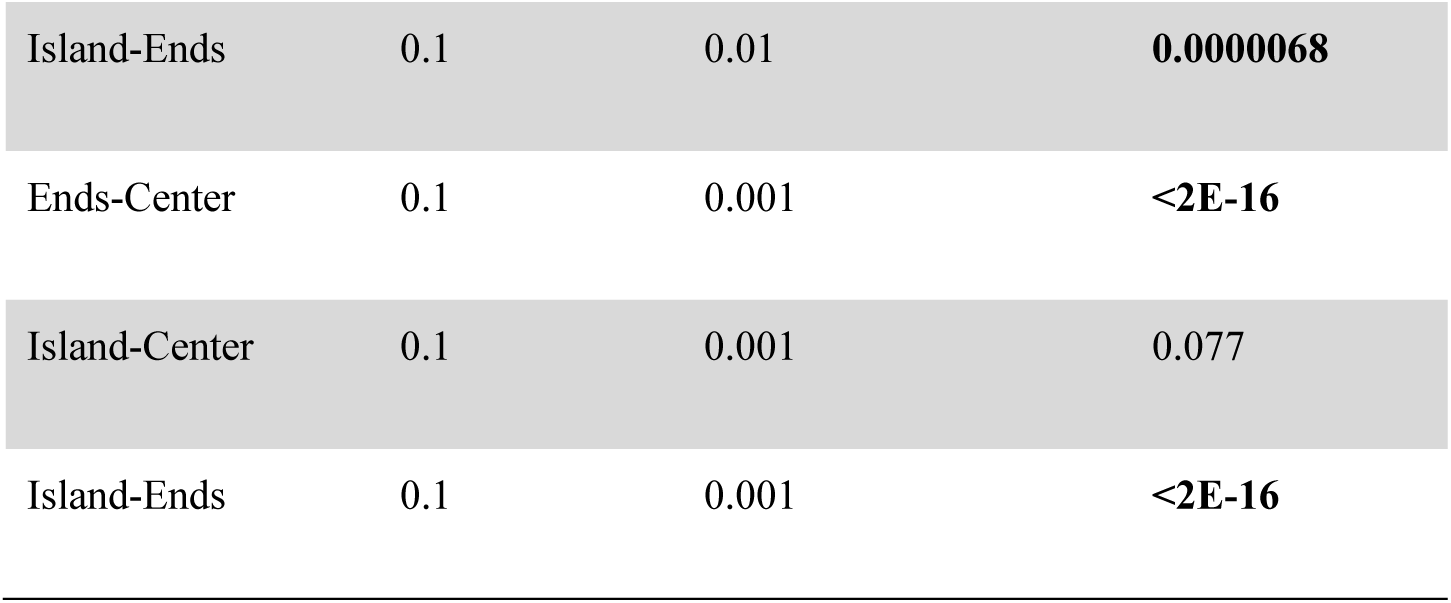
Pairwise Wilcoxon test results for comparisons of T_D_ / *ND* of 10, 5, 2, 1, 0.5, and 0.1. Significant (*p* < 0.05) results are bolded.

**Table S2.** Wilcoxon pairwise test comparing (T_MRCA_ – T_D_) / T_D_ of center and end demes for different rates of migration and ratios of T_D_ / *ND*. Bolded *p*-values indicate *p* < 0.05.

| Ratio | Migration rate | p-value |
| --- | --- | --- |
| 50 | 0.1 | 0.385 |
| 50 | 0.01 | **0.000748** |
| 50 | 0.001 | **2.00E-16** |
| 25 | 0.1 | **0.00282** |
| 25 | 0.01 | **3.55E-09** |
| 25 | 0.001 | **2.00E-16** |
| 10 | 0.1 | **0.0118** |
| 10 | 0.01 | **1.24E-08** |
| 10 | 0.001 | **2.00E-16** |
| 5 | 0.1 | 0.744 |
| 5 | 0.01 | 0.0528 |
| 5 | 0.001 | **2.00E-16** |
| 1 | 0.1 | 0.0672 |
| 1 | 0.01 | **2.00E-16** |
| 1 | 0.001 | **2.00E-16** |

**Table S3.** Table of estimated divergence-times in SNAPP.

| $T_D / ND$ | Deme sampled | Migration rate | True Age ($T_D$) | Expected Estimate ($eT_D$) | Actual Estimate ($T_{MRCA}$) | $T_{MRCA} - eT_D$ | $(T_{MRCA} - eT_D) / eT_D$ |
| --- | --- | --- | --- | --- | --- | --- | --- |
| 50 | End | 0.1 | 50000 | 52000 | 32929 | -19071 | -0.36675 |
| 50 | End | 0.01 | 50000 | 52000 | 36330 | -15670 | -0.301346154 |
| 50 | End | 0.001 | 50000 | 52000 | 40028 | -11972 | -0.230230769 |
| 50 | Center | 0.1 | 50000 | 52000 | 46270 | -5730 | -0.110192308 |
| 50 | Center | 0.01 | 50000 | 52000 | 54301 | 2301 | 0.04425 |
| 50 | Center | 0.001 | 50000 | 52000 | 49004 | -2996 | -0.057615385 |
| 25 | End | 0.1 | 25000 | 27000 | 25893 | -1107 | -0.041 |
| 25 | End | 0.01 | 25000 | 27000 | 26492 | -508 | -0.018814815 |
| 25 | End | 0.001 | 25000 | 27000 | 48430 | 21430 | 0.793703704 |
| 25 | Center | 0.1 | 25000 | 27000 | 22264 | -4736 | -0.175407407 |
| 25 | Center | 0.01 | 25000 | 27000 | 25488 | -1512 | -0.056 |
| 25 | Center | 0.001 | 25000 | 27000 | 26147 | -853 | -0.031592593 |
| 10 | End | 0.1 | 10000 | 12000 | 12439 | 439 | 0.036583333 |
| 10 | End | 0.01 | 10000 | 12000 | 14085 | 2085 | 0.17375 |
| 10 | End | 0.001 | 10000 | 12000 | 28629 | 16629 | 1.38575 |
| 10 | Center | 0.1 | 10000 | 12000 | 13234 | 1234 | 0.102833333 |
| 10 | Center | 0.01 | 10000 | 12000 | 13040 | 1040 | 0.086666667 |
| 10 | Center | 0.001 | 10000 | 12000 | 21782 | 9782 | 0.815166667 |
| 5 | End | 0.1 | 5000 | 7000 | 10618 | 3618 | 0.516857143 |
| 5 | End | 0.01 | 5000 | 7000 | 12730 | 5730 | 0.818571429 |
| 5 | End | 0.001 | 5000 | 7000 | 19936 | 12936 | 1.848 |
| 5 | Center | 0.1 | 5000 | 7000 | 11144 | 4144 | 0.592 |
| 5 | Center | 0.01 | 5000 | 7000 | 10595 | 3595 | 0.513571429 |
| 5 | Center | 0.001 | 5000 | 7000 | 11697 | 4697 | 0.671 |
| 1 | End | 0.1 | 1000 | 3000 | 1512.9 | -1487.1 | -0.4957 |
| 1 | End | 0.01 | 1000 | 3000 | 2700 | -300 | -0.1 |
| 1 | End | 0.001 | 1000 | 3000 | 23845 | 20845 | 6.948333333 |
| 1 | Center | 0.1 | 1000 | 3000 | 1513.2 | -1486.8 | -0.4956 |
| 1 | Center | 0.01 | 1000 | 3000 | 2649.1 | -350.9 | -0.116966667 |
| 1 | Center | 0.001 | 1000 | 3000 | 1647.2 | -1352.8 | -0.450933333 |

**Figure S1.**
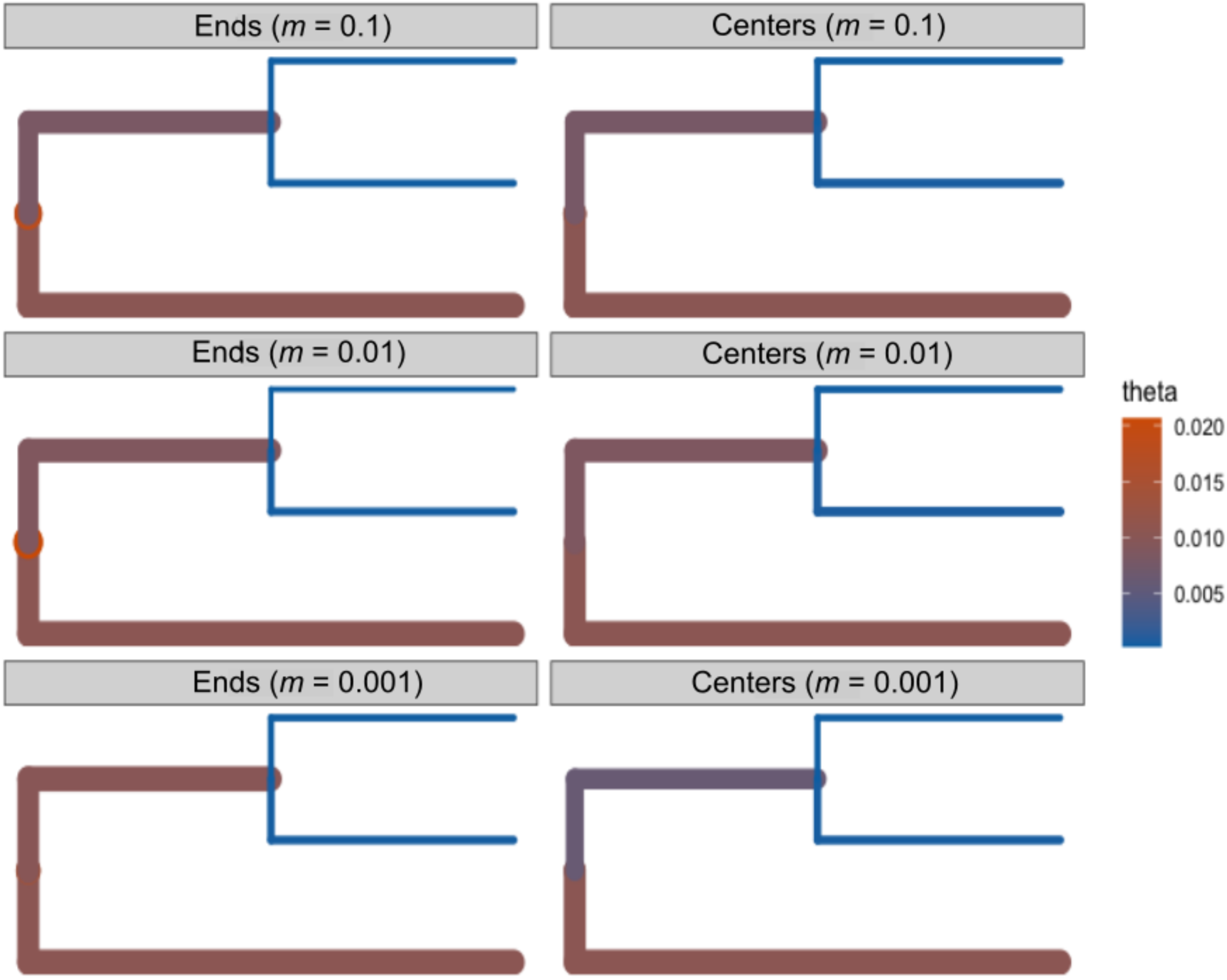
Estimates of *θ* in SNAPP for T_D_ / *ND* = 50. Branch widths are proportional to the estimated *θ*.

**Figure S2.**
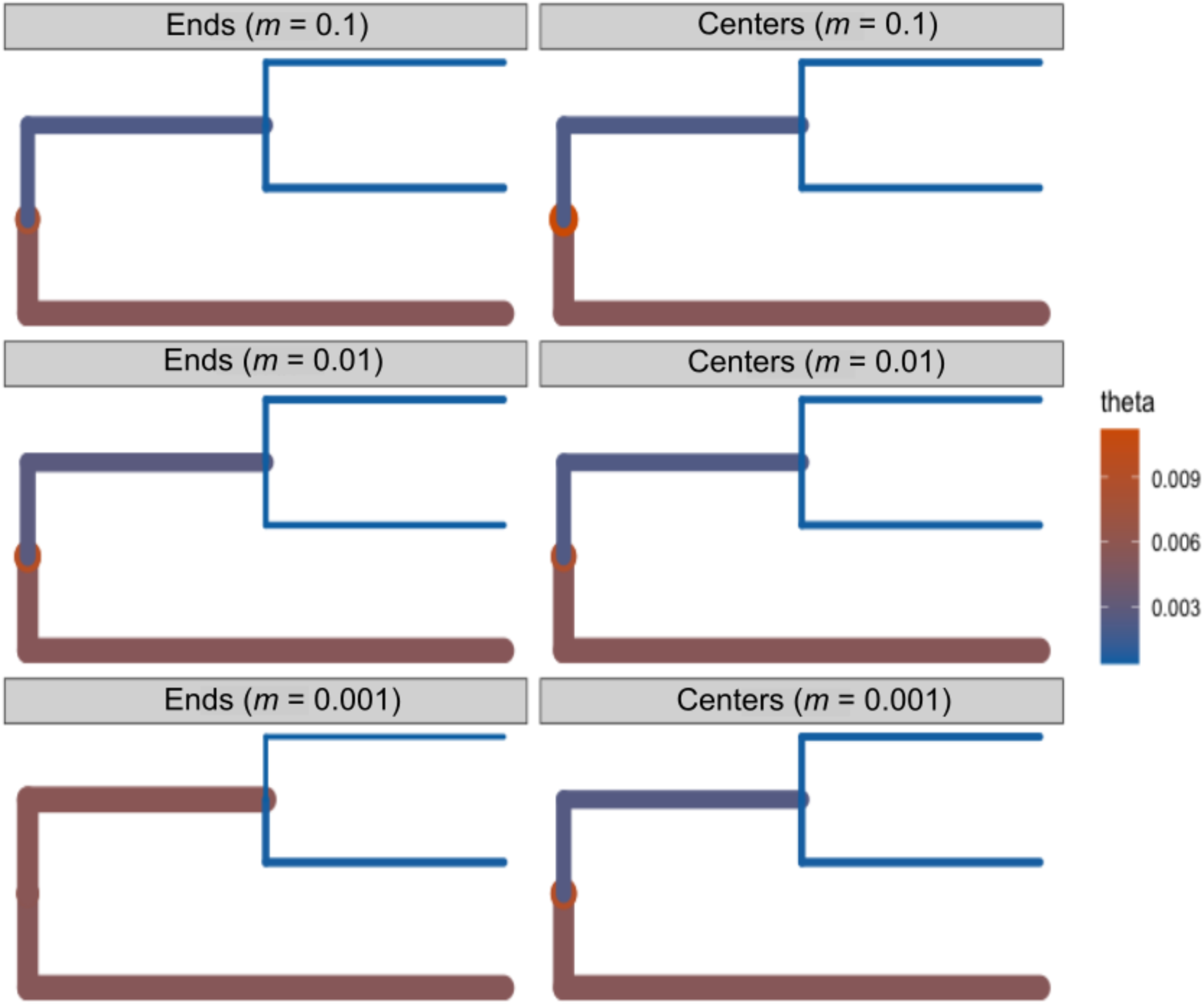
Estimates of *θ* in SNAPP for T_D_ / *ND* = 25. Branch widths are proportional to the estimated *θ*.

**Figure S3.**
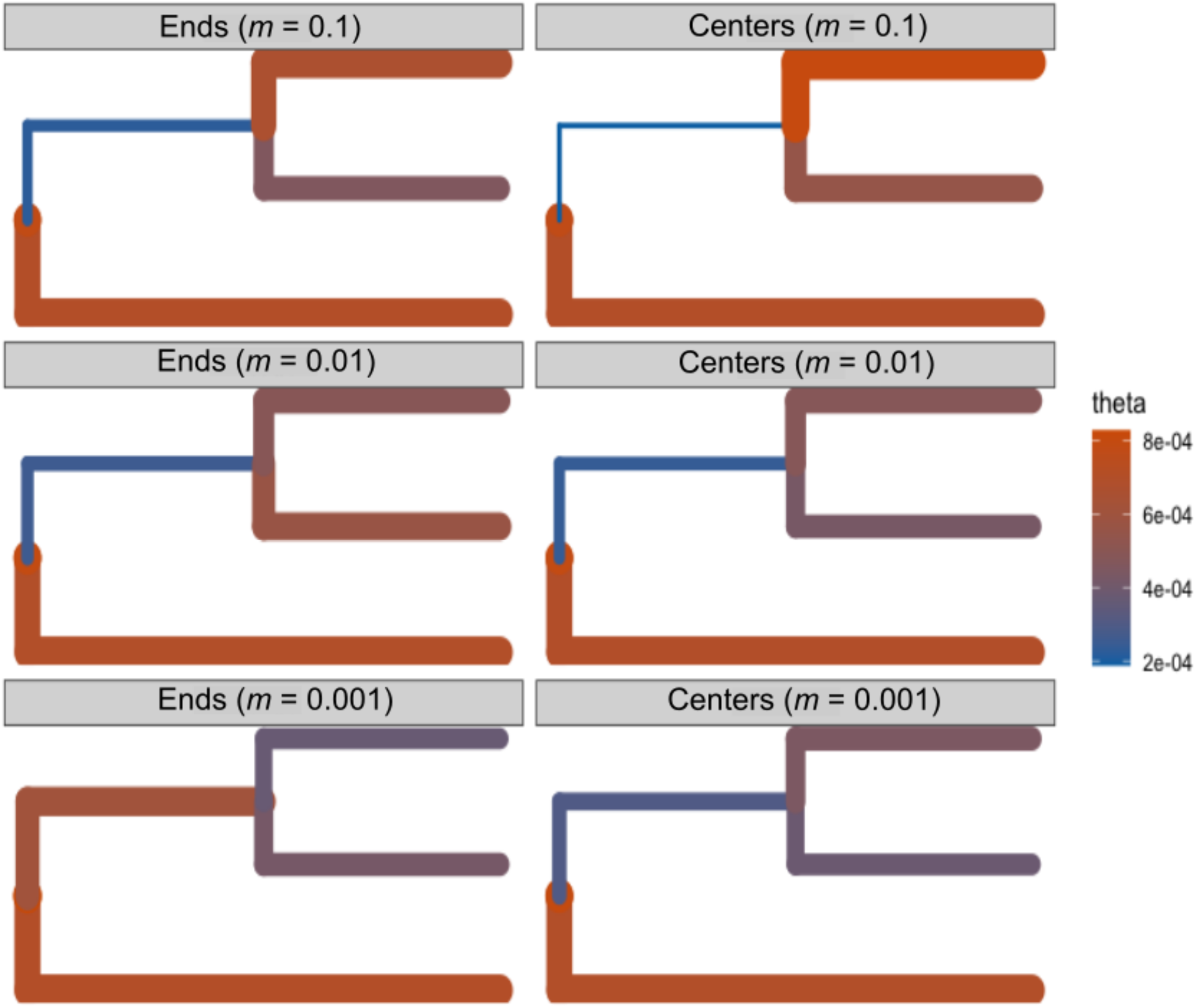
Estimates of *θ* in SNAPP for T_D_ / *ND* = 5. Branch widths are proportional to the estimated *θ*.

**Figure S4.**
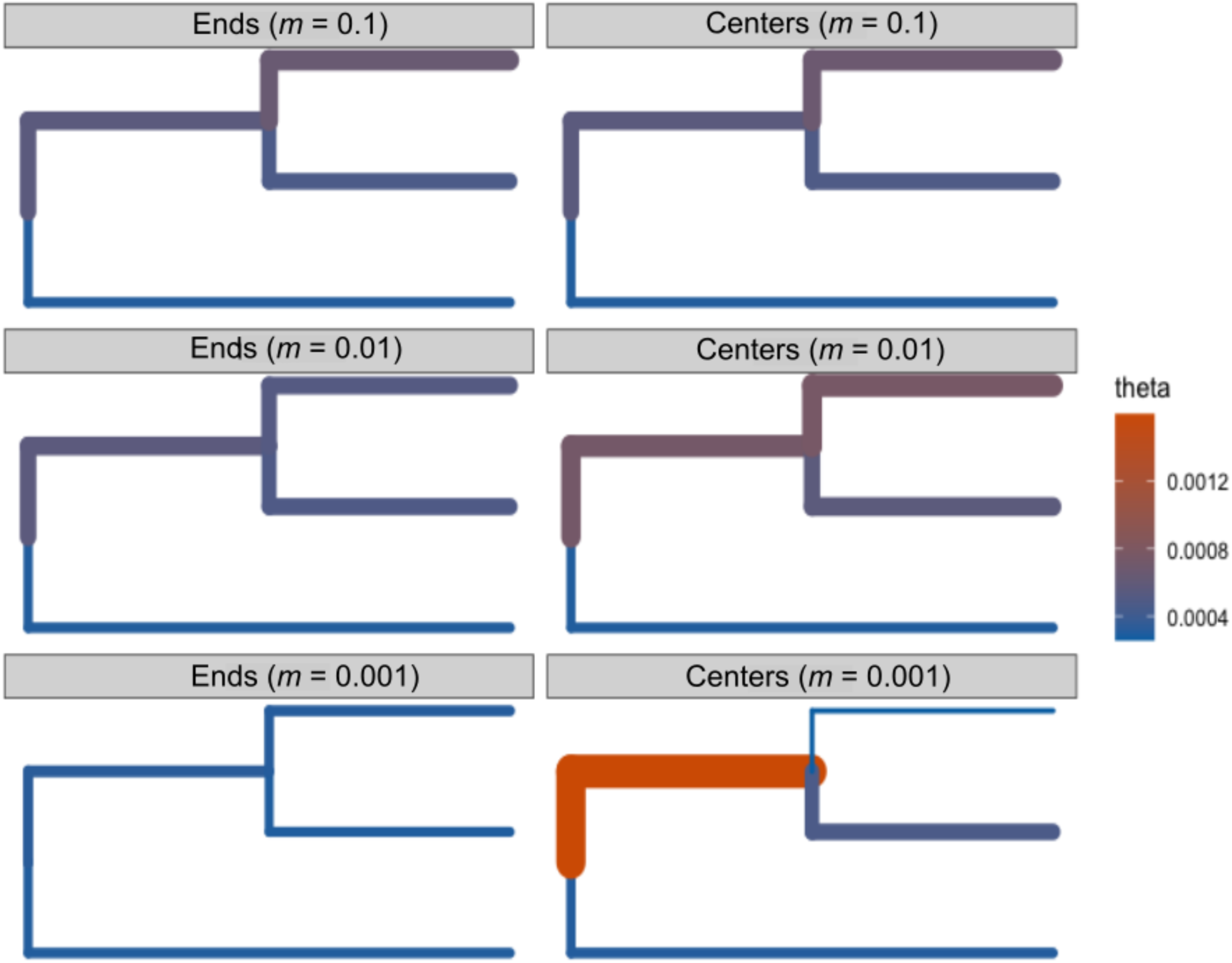
Estimates of *θ* in SNAPP for T_D_ / *ND* = 1. Branch widths are proportional to the estimated *θ*.

**Figure S5.**
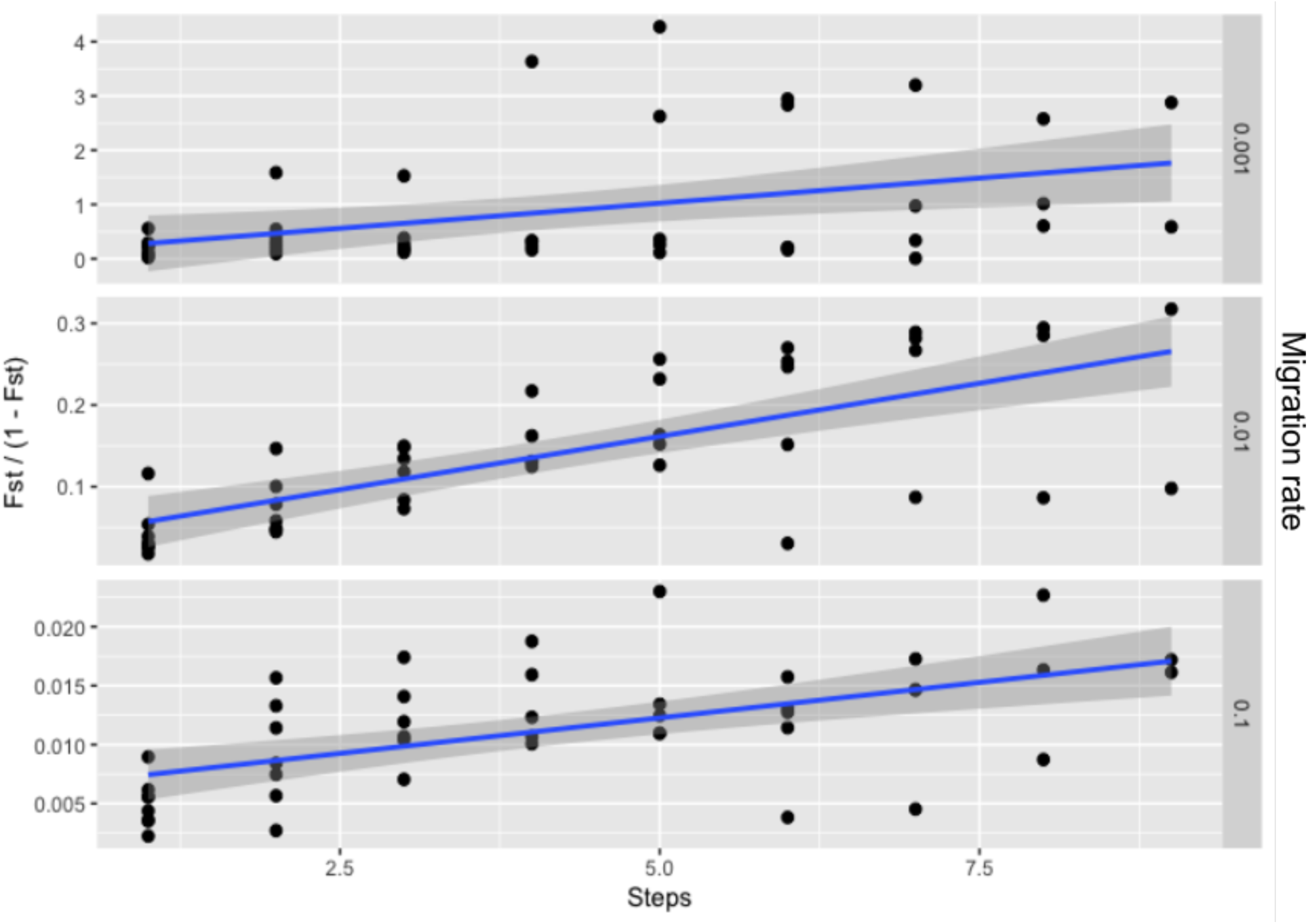
Isolation-by-distance plots for three migration rates in the ancestral population. Pairwise *F*_ST_ was calculated between each deme in the ancestral population prior to the split to verify that a pattern of IBD had occurred. “Steps” are the distance from deme *i* to deme *j*, where neighboring demes are 1 step apart. Note that the y-axis differs between panels. All slopes were significant (*p* < 0.05).

